# A SARS-CoV-2 BioID-based virus-host membrane protein interactome and virus peptide compendium: new proteomics resources for COVID-19 research

**DOI:** 10.1101/2020.08.28.269175

**Authors:** Jonathan R. St-Germain, Audrey Astori, Payman Samavarchi-Tehrani, Hala Abdouni, Vinitha Macwan, Dae-Kyum Kim, Jennifer J. Knapp, Frederick P. Roth, Anne-Claude Gingras, Brian Raught

**Affiliations:** Princess Margaret Cancer Centre, University Health Network, 101 College St., Toronto, ON, Canada; Lunenfeld-Tanenbaum Research Institute, Sinai Health System, Toronto, ON, Canada; Department of Molecular Genetics, University of Toronto, Toronto, ON, Canada; Department of Computer Science, University of Toronto, Toronto, ON, Canada; Department of Medical Biophysics, University of Toronto, Toronto, ON, Canada

**Author notes:** these authors contributed equally (Audrey Astori, Payman Samavarchi-Tehrani).

## Abstract

Key steps of viral replication take place at host cell membranes, but the detection of membrane-associated protein-protein interactions using standard affinity-based approaches (e.g. immunoprecipitation coupled with mass spectrometry, IP-MS) is challenging. To learn more about SARS-CoV-2 - host protein interactions that take place at membranes, we utilized a complementary technique, proximity-dependent biotin labeling (BioID). This approach uncovered a virus-host topology network comprising 3566 proximity interactions amongst 1010 host proteins, highlighting extensive virus protein crosstalk with: (i) host protein folding and modification machinery; (ii) membrane-bound vesicles and organelles, and; (iii) lipid trafficking pathways and ER-organelle membrane contact sites. The design and implementation of sensitive mass spectrometric approaches for the analysis of complex biological samples is also important for both clinical and basic research proteomics focused on the study of COVID-19. To this end, we conducted a mass spectrometry-based characterization of the SARS-CoV-2 virion and infected cell lysates, identifying 189 unique high-confidence virus tryptic peptides derived from 17 different virus proteins, to create a high quality resource for use in targeted proteomics approaches. Together, these datasets comprise a valuable resource for MS-based SARS-CoV-2 research, and identify novel virus-host protein interactions that could be targeted in COVID-19 therapeutics.

## Introduction

The SARS-CoV-2 ∼30 kb positive strand RNA genome (Genbank MN908947.3^1, 2^) contains two large open reading frames (ORF1a and ORF1ab) encoding polyproteins that are cleaved by viral proteases into ∼16 non-structural proteins (NSPs). 13 smaller 3’ ORFs encode the primary structural proteins of the virus, spike (S), nucleocapsid (N), membrane (M) and envelope (E), along with nine additional polypeptides of poorly understood function (**Fig 1A**).

**Figure 1.**
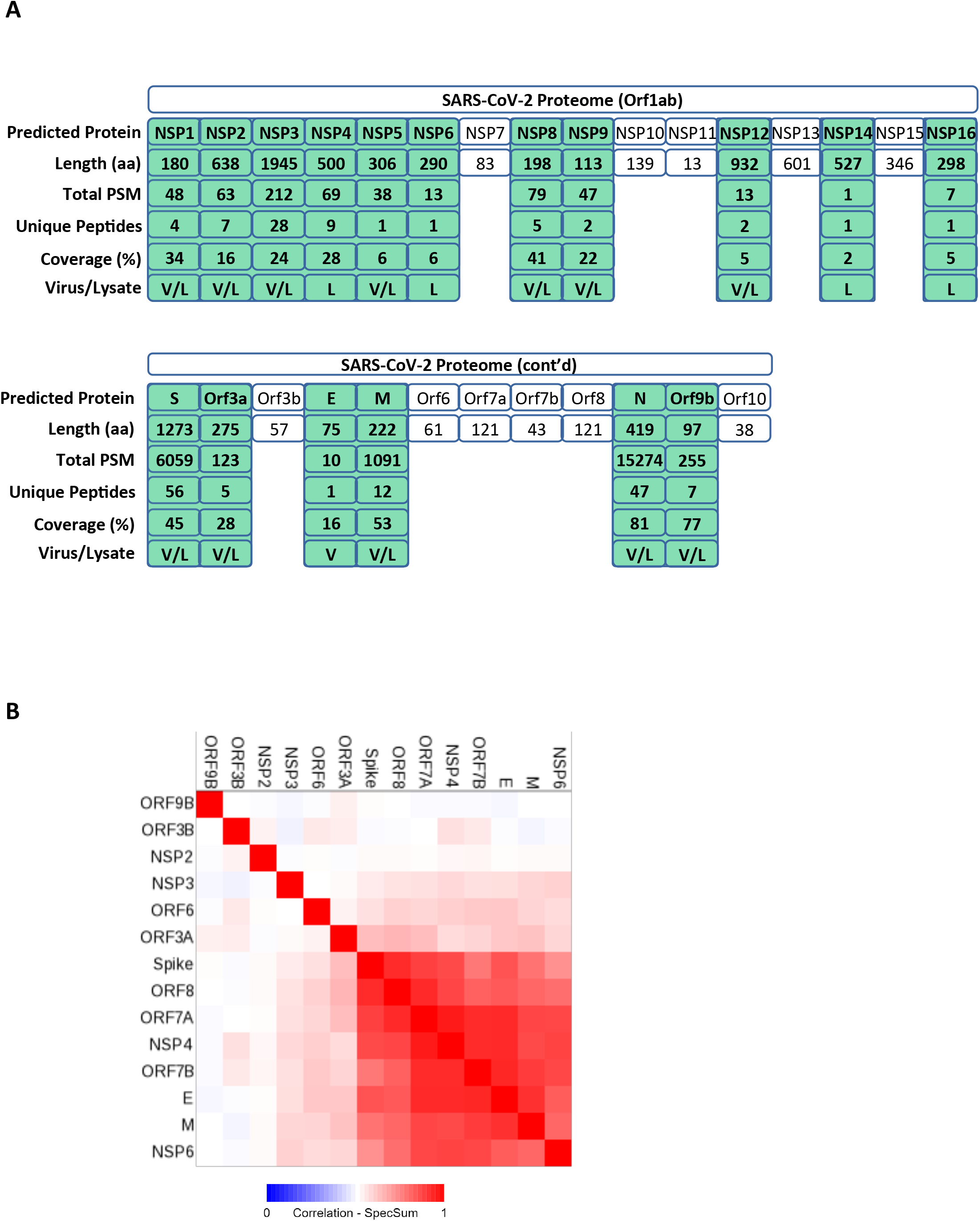
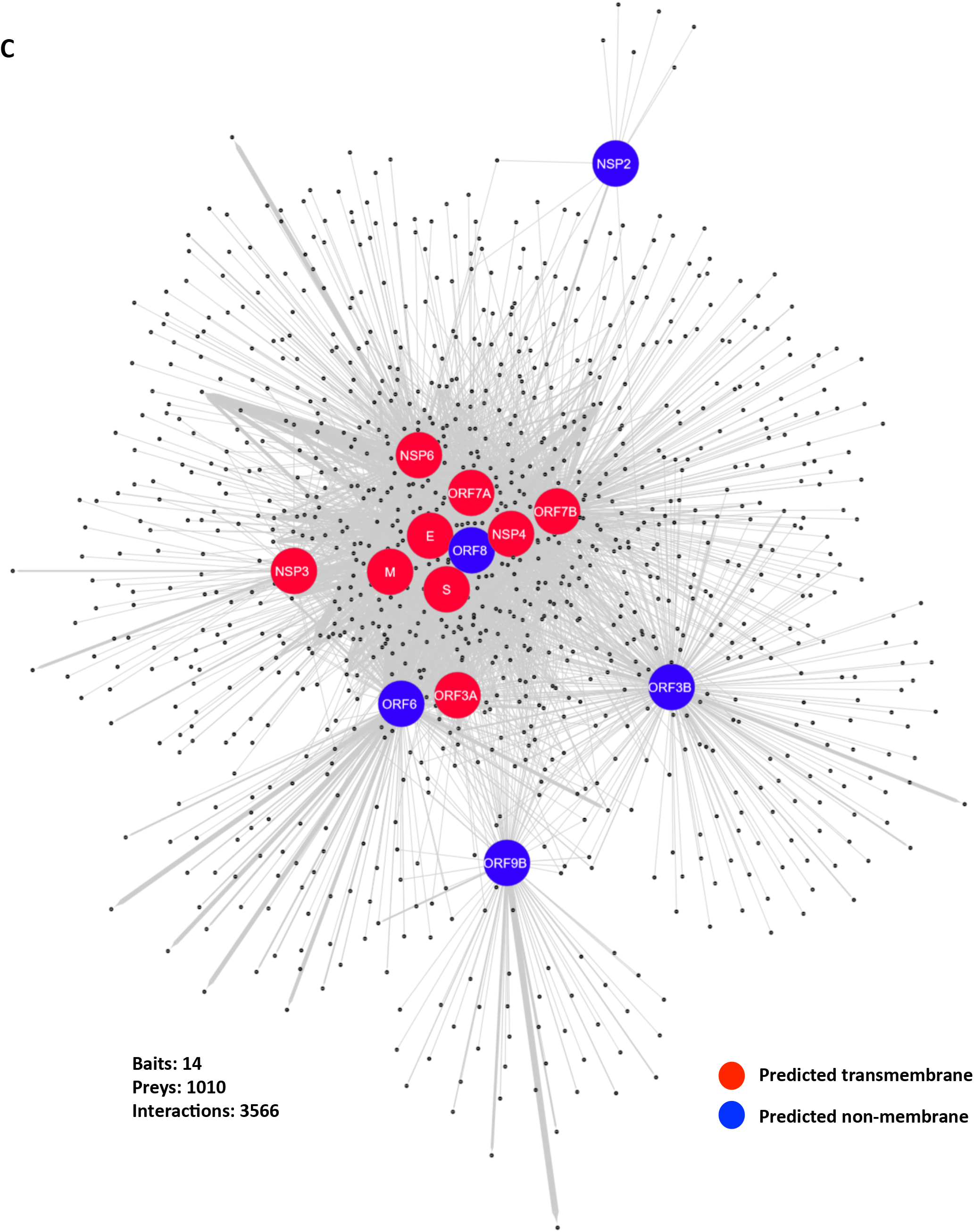
SARS-CoV-2 - host proximity interaction mapping. (A) Polypeptides encoded by the SARS-CoV-2 (+)RNA genome, and detected in DDA mass spectrometry, as indicated. (B) Bait-bait clustering. Interactomes were clustered according to Jaccard similarity index, using the ProHits-viz^17^ (prohits-viz.lunenfeld.ca) correlation analysis tool. (C) Self-organized prefuse force-directed topology map (based on peptide count sum of four mass spectrometry runs) for the SARS-CoV-2 BioID interactome. Baits are indicated with large Red (one or more predicted transmembrane helices) or Blue (no predicted transmembrane regions) circles. Prey are indicated by smaller black dots. Edge width is proportional to spectral counts.

To enable proteomics-based approaches for the analysis of complex biological samples, we analyzed both SARS-CoV-2-infected cell lysates and mature virions, generating a high confidence virus peptide spectrum compendium. This dataset can be used e.g. for the selection of virus peptides for use in targeted proteomics approaches (e.g. the identification of viral peptides in human clinical samples), or for the generation of peptide spectral libraries for increased sensitivity of detection.

Two SARS-CoV-2 - host protein-protein interaction (PPI) mapping efforts have utilized immunoprecipitation coupled with mass spectrometry (IP-MS) of epitope-tagged viral proteins to identify >1000 putative virus-host PPIs in HEK293T^3^ and A549^4^ cells. While extremely powerful for identifying stable, soluble protein complexes, IP-MS approaches are not optimal for the capture of weak or transient PPIs, or for the detection of PPIs that take place at poorly soluble intracellular locations such as membranes, where key steps in viral replication occur.

To better understand how host cell functions are hijacked and subverted by SARS-CoV-2 proteins, we used proximity-dependent biotinylation (BioID^5^) as a complementary approach to map virus-host protein proximity interactions in live human cells. This dataset provides a valuable resource for better understanding SARS-CoV-2 pathogenesis, and identifies numerous previously undescribed virus-host PPIs that could represent attractive targets for therapeutic intervention.

## Results

### A SARS-CoV-2 peptide compendium

To identify high quality tryptic virus peptides for use in targeted proteomics analyses, data-dependent acquisition (DDA) mass spectrometry was conducted on the mature SARS-CoV-2 virion. The Toronto SB3 virus strain^6^ was cultured in VeroE6 cells (MOI 0.1), and culture media was collected 48 hrs post-infection. Virus was concentrated by centrifugation, inactivated by detergent and subjected to tryptic proteolysis. The resulting viral tryptic peptides were identified using nanoflow liquid chromatography - tandem mass spectrometry (LC-MS/MS; **Fig 1A, Supp Table 1**). Multiple unique peptides of the structural proteins N, S, and M (but only a single unique peptide for E) were detected in this analysis. Lower numbers of peptides derived from several NSPs were also reproducibly detected in mature virion preparations.

SARS-CoV-2 proteins produced in the context of an infection were similarly analyzed. Lysates from infected VeroE6 cells (MOI 0.1, 48 hrs post-infection) were inactivated with detergent, subjected to trypsin proteolysis and analyzed by DDA LC-MS/MS. In this case, unique peptides derived from 16 different virus proteins were detected (**Fig1A, Supp Table 1**; all virus MS data available at massive.ucsd.edu, accession #MSV000086017).

Together, these data confirm and expand upon previous proteomic analyses of SARS-CoV-2 virions, infected cells^4, 7-11^ and patient samples^12-14^, and provide a library of high quality virus peptide spectra covering 17 virus proteins that can be used for the creation of peptide spectral libraries and targeted proteomics approaches.

### A SARS-CoV-2 - host protein proximity interactome

Based on standard transcript mapping algorithms and conservation with ORFs in other coronaviruses^1, 2^, we created a SARS-CoV-2 open reading frame (ORF) vector set^15^ (**Fig 1A**). Nine SARS-CoV-2 proteins are predicted to have one or more transmembrane domains (S, E, M, NSP3, NSP4, NSP6, ORF3A, ORF7A and ORF7B). To better characterize SARS-CoV-2 - host membrane-associated PPIs, these virus ORFs (along with the remaining poorly understood open reading frames ORF3B, ORF6, ORF8 and ORF9B) were fused in-frame with an N-terminal BirA* (R118G) coding sequence, and the resulting fusion proteins individually expressed in HEK 293 Flp-In T-REx cells. Using these cells, a virus-host PPI landscape was characterized using BioID (as in^16^; **Supp Table 2**). Applying a Bayesian false discovery rate of ≤1%, 3566 high confidence proximity interactions were identified with 1010 unique human proteins (all raw data available at massive.ucsd.edu, accession #MSV000086006). 412 prey polypeptides were detected as high confidence interactors for a single SARS-CoV-2 bait protein in this analysis, underscoring the high degree of specificity in this virus-host proximity interaction map.

Bait-bait correlation analysis (**Fig 1B**), based on similarity between interactomes (Jaccard Index analysis conducted in ProHits-viz^17^) revealed high levels of correspondence between the S (Spike), E, M, NSP4, NSP6, ORF7A, ORF7B, and ORF8 bait proteins. The NSP2, NSP3, ORF3A, ORF3B, ORF6, and ORF9B interactomes shared a lower degree of similarity with the other bait proteins in this set. Bait proteins with one or more predicted transmembrane domains thus largely clustered together, with the exception of ORF3A (which clusters outside the main group of putative membrane baits, even though it is predicted to possess three transmembrane helices), and ORF8 (which clusters with the putative membrane baits, but has no predicted transmembrane domain itself).

A self-organized force-directed bait-prey topology map was next generated, in which map location is determined by the number and abundance (i.e. total peptide counts) of host cell interactors (**Fig 1C**). This approach similarly clustered all of the baits with one or more predicted transmembrane helices, along with ORF8, in a dense “core” region of the map, indicating that these bait proteins share a large proportion of common interactors. NSP3 and ORF3A occupy regions at the edge of this dense region of the map, indicating a lower number of shared interactors with the other membrane proteins. NSP2, ORF3B and ORF9B occupy peripheral regions of the map, indicating that they share far fewer interactors with the rest of the baits analyzed here. Interestingly, ORF6 occupies a region of the map near ORF3A (these two baits were also clustered near each other in the bait-bait analysis, above). Consistent with this location, more than half of the ORF3A interactome (39 of 73 proteins) is also present in the larger ORF6 interactome (217 proteins). This overlapping group of interactors is enriched in plasma membrane (PM) and ER proteins. Based on this observation, it will be interesting to explore similarities in ORF3A and ORF6 function.

### SARS-CoV-2 proteins display extensive host membrane protein interactions

As a whole, the virus-host interactome is significantly enriched in proteins associated with the endoplasmic reticulum (ER)/nuclear, Golgi and plasma membranes, and ER-Golgi trafficking vesicles (**Table 1, Supp Table 3**). Enriched biological functions include ER-Golgi vesicle-mediated transport, response to ER stress, and lipid biosynthesis. Significantly enriched molecular functions (i.e. enriched protein domains or motifs with known functions) include SNARE binding, lipid transporter activity, and sterol binding activities (**Table 1, Supp Table 3**).

**Table 1.**
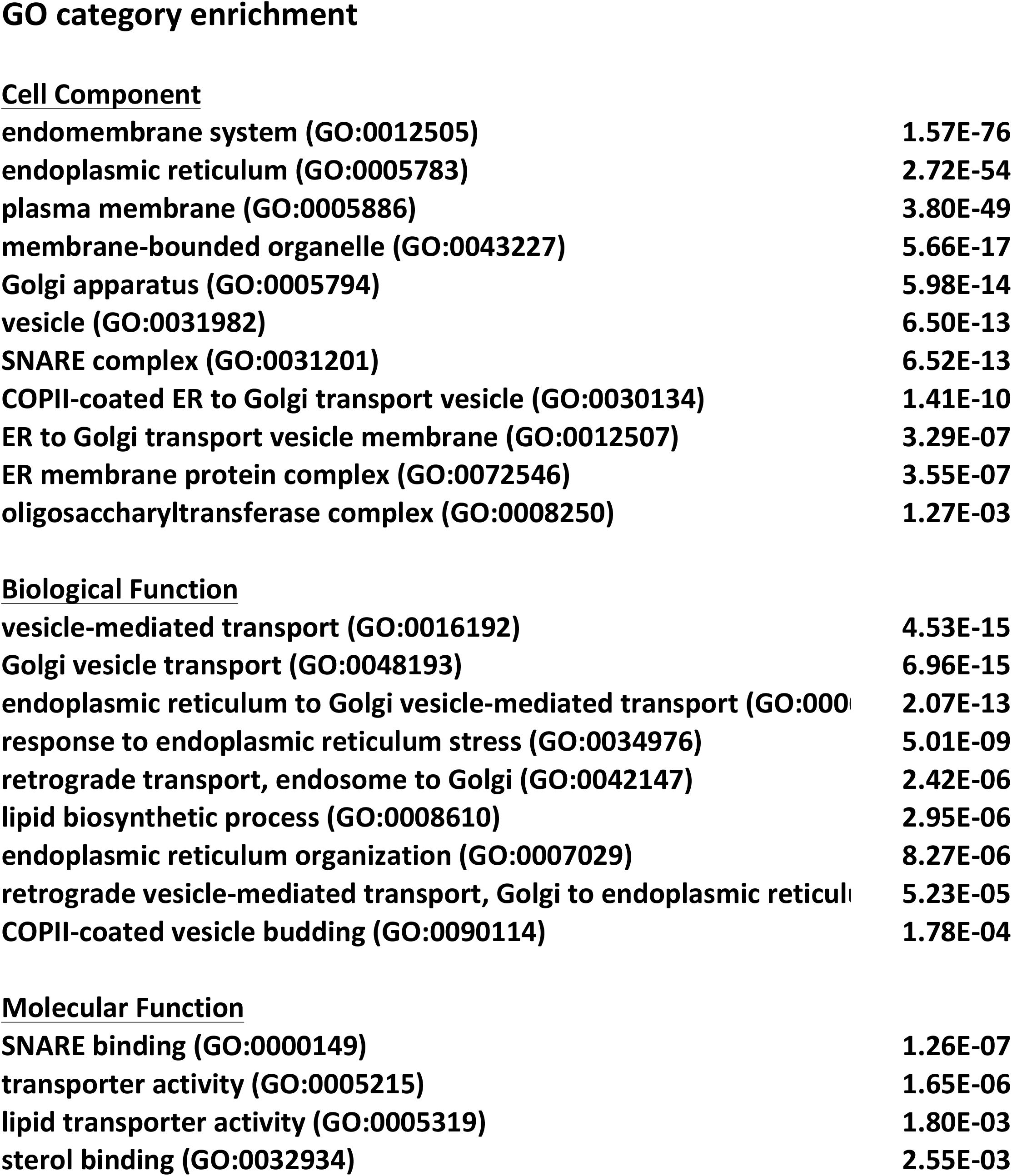
Significantly enriched gene ontology (GO) categories (selected), and associated FDR, for the complete BioID dataset.

|  |  |
| --- | --- |
| endomembrane system (GO:0012505) | 1.57E-76 |
| endoplasmic reticulum (GO:0005783) | 2.72E-54 |
| plasma membrane (GO:0005886) | 3.80E-49 |
| membrane-bounded organelle (GO:0043227) | 5.66E-17 |
| Golgi apparatus (GO:0005794) | 5.98E-14 |
| vesicle (GO:0031982) | 6.50E-13 |
| SNARE complex (GO:0031201) | 6.52E-13 |
| COPII-coated ER to Golgi transport vesicle (GO:0030134) | 1.41E-10 |
| ER to Golgi transport vesicle membrane (GO:0012507) | 3.29E-07 |
| ER membrane protein complex (GO:0072546) | 3.55E-07 |
| oligosaccharyltransferase complex (GO:0008250) | 1.27E-03 |
| **Biological Function** |  |
| vesicle-mediated transport (GO:0016192) | 4.53E-15 |
| Golgi vesicle transport (GO:0048193) | 6.96E-15 |
| endoplasmic reticulum to Golgi vesicle-mediated transport (GO:0006691) | 2.07E-13 |
| response to endoplasmic reticulum stress (GO:0034976) | 5.01E-09 |
| retrograde transport, endosome to Golgi (GO:0042147) | 2.42E-06 |
| lipid biosynthetic process (GO:0008610) | 2.95E-06 |
| endoplasmic reticulum organization (GO:0007029) | 8.27E-06 |
| retrograde vesicle-mediated transport, Golgi to endoplasmic reticulum (GO:0006692) | 5.23E-05 |
| COPII-coated vesicle budding (GO:0090114) | 1.78E-04 |
| **Molecular Function** |  |
| SNARE binding (GO:0000149) | 1.26E-07 |
| transporter activity (GO:0005215) | 1.65E-06 |
| lipid transporter activity (GO:0005319) | 1.80E-03 |
| sterol binding (GO:0032934) | 2.55E-03 |

Multiple SARS-CoV-2 bait proteins interact with the ER signal peptidase complex (SEC11A, SEC11C, SPCS1-3), the ER membrane complex (TMEM85, TMEM111, MMGT, FAM158A, KIAA0090, TTC35, COX4NB), the intramembrane ER chaperone CCDC47^18^, and the oligosaccharyltransferase (OST) complex (RPN1, RPN2, DDOST, MAGT1, STT3B, TUSC3). This suite of ER-based host proteins is thus likely to be required for efficient SARS-CoV-2 polypeptide folding, post-translational modications, membrane localization and virion assembly.

Multiple SARS-CoV-2 polypeptides also interact with intracellular vesicle trafficking proteins (SNX19, RAB7A, RAB23 and others), and protein components of membrane-bound organelles, including lipid droplets (AUP1, LPCAT, LPGAT, and MBOAT proteins), peroxisomes (ABCD3/PMP70, ACBD5) and mitochondria (FAM82A2, TMEM173, CYB5A, CYB5B). The virus protein interactome is also notably enriched in membrane contact site (MCS) components. Direct inter-organelle sterol and lipid transfer is executed at MCSs, where protein-protein interactions tether two organellar membranes together to create microdomains suitable for the direct exchange of lipids and sterols^19^. Key to this process are the ER membrane tethering proteins VAPA, VAPB and MOSPD2. SARS-CoV-2 proteins display extensive interactions with all of the ER tethers, a variety of MCS components, and almost all known ER-organelle tethering proteins, including: ESYT1, ESYT2, STIM1, STIM2, C2CD2L and GRAMD1A/B/C plasma membrane-ER MCS proteins; the peroxisome-ER tether, ACBD5; the mitochondria-ER MCS proteins PDZD8, VPS13A, INF2, FAM82A2/RMDN3, ITPR1 and ITPR3; the Golgi-ER tether PLEKHA8/FAPP2; and the lipid droplet-ER MCS proteins RAB3GAP1, RAB3GAP2, STX18, USE1, NBAS and RINT1 (**Figure 2**; **Supp Table 2**). The interactome is also enriched with a number of cholesterol/lipid transfer proteins, including multiple members of the oxysterol binding protein family; OSBP, OSBPL2, OSBPL3, OSBPL8, OSBPL9, OSBPL10 and OSBPL11.

**Figure 2.**
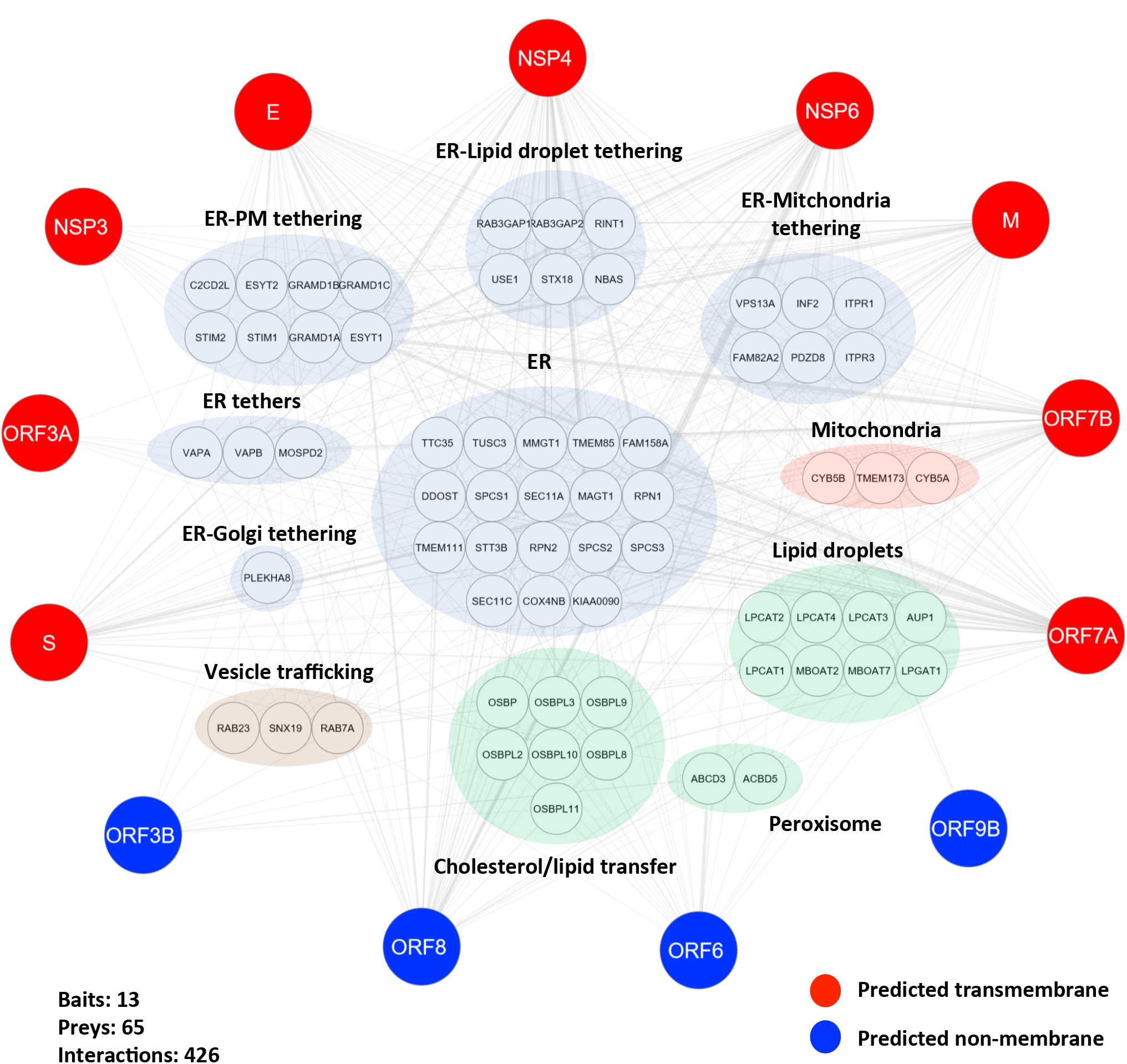
SARS-CoV-2 proteins display physical and functional links with membrane contact site components and lipid/sterol transport proteins. Virus bait protein proximity interactions with host membrane protein functional groups are highlighted.

All positive-strand RNA viruses effect a dramatic rearrangement of host cell membranes to create replication organelles (ROs, also referred to as double membrane vesicles, or DMVs), membrane-bound “virus factories” that scaffold and protect the viral replicative machinery from host defenses^20^. (+)RNA viruses can also manipulate the host cell lipid and cholesterol trafficking machinery to produce RO membranes with much higher phosphatidylinositol 4-phosphate (PI(4)P) and cholesterol content than the host ER/Golgi membranes from which they are derived. To this end, many enteroviruses exploit the PI(4)P-driven cholesterol transport system normally used at ER MCSs, via recruitment of OSBP/OSBPL proteins to the RO membrane^21-24^. The abundant interactions of SARS-CoV-2 proteins with host MCS components and lipid/cholesterol transfer proteins strongly suggests that this virus manipulates lipid trafficking functions in a similar manner. Disruption of PI(4)P-driven cholesterol transport or OSBP/L function blocks the replication of several virus types, including HCV, poliovirus, EMCV and enteroviruses^20-22, 24, 25^. The human MCS lipid transfer system may thus represent an interesting drug target for COVID-19 treatment.

Even amongst those viral proteins that appear to localize exclusively to the ER-Golgi-PM endomembrane membrane system, specificity in virus-host interactomes was observed, likely reflecting preferences for interactions with different subsets of membrane proteins and/or localization to unique membrane lipid nanodomains. For example, both the ORF7A and ORF7B interactomes are enriched in PM solute channels, but ORF7A appears to interact uniquely with the anion exchanger SLC4A2, the taurine transporter SLC6A6, and the glycine transporter SLC6A9, while ORF7B interacts specifically with the amino acid transporter SLC1A5, the sulfate transporter SLC26A11, and the divalent metal transporter SLC39A14.

Autophagy is an important part of the innate immune response, effecting the elimination of intracellular pathogens such as viruses (virophagy), and delivering them to the lysosome, which processes pathogen components for antigen presentation^26, 27^. Many viruses have thus evolved strategies to inhibit the host autophagic machinery. Notably, however, (+)RNA viruses appear to be dependent on autophagic function for efficient replication, and hijack components of the autophagic machinery for use in membrane re-organization and the creation of ROs^28^. Consistent with these observations, our data highlight multiple interactions amongst SARS-CoV-2 proteins and the ER-phagy receptors FAM134B, TEX2644 and SEC24C. A number of virus protein interactions were also detected with components of the UFMylation system (DDRGK1, CDK5RAP3, UFL1 and UFSP2), which was recently shown to play a key role in ER-phagy^10^, highlighting interesting links between specific autophagy pathways and SARS-CoV-2.

### Crosstalk with other organelles and host protein machines

Consistent with that observed in IP-MS analyses - and unlike any of the other baits analyzed here - the SARS-CoV-2 ORF9B interactome is significantly enriched in mitochondrial proteins (**Supp Table 2, Supp Table 3**). Amongst the high confidence ORF9B interactors is the mitochondrial antiviral signaling protein (MAVS), which acts as a hub for cell-based innate immune signaling. The cellular pattern recognition receptors (PRRs) detect pathogen-associated molecular patterns (PAMPs, e.g. viral RNAs). PAMP-bound PRRs interact with MAVS, which activates the NF-kB and Type I interferon signaling pathways^29^. Many different viruses block the host antiviral response by interfering with MAVS signaling. This may be accomplished by e.g. direct MAVS cleavage by viral proteases (a strategy used by HAV, HCV and coxsackievirus B3), or via 26S proteasome-mediated degradation, a strategy used by SARS-CoV ORF9B^30^, which recruits the HECT E3 ligase ITCH/AIP4 to effect MAVS ubiquitylation. We did not detect any ubiquitin E3 ligases in the ORF9B BioID interactome, but consistent with a recent report indicating that SARS-CoV-2 ORF9B binds directly to TOMM70^31^ to block MAVS-mediated IFN signaling, we detected TOMM70 as a major component of the ORF9B interactome.

The SARS-CoV-2 ORF6 interactome is uniquely enriched in nuclear pore complex (NPC) components (**Supp Table 2, Supp Table 3**). SARS-CoV ORF6 was shown to inhibit NPC-mediated transport by tethering the importin proteins KPNA2 and KPNB2/TNPO1 to ER/Golgi membranes^32^, which effectively blocks the import of immune signaling proteins such as STAT1 into the nucleus. SARS-CoV-2 ORF6 shares 69% identity at the amino acid level with its SARS-CoV counterpart, and similarly displays potent immune repressor function^33^. It will be interesting to detemine if the SARS-CoV2 ORF6 interaction with NPC components leads to a similar disruption of immune signaling and nuclear transport.

ORF3B interacts specifically with LAMTOR1 and LAMTOR2, components of the Ragulator complex, which is localized to the lysosomal membrane and regulates the mechanistic target of rapamycin complex 1 (mTORC1). mTOR signaling is inactivated by amino acid starvation and other types of stress, to inhibit cap-dependent translation and upregulate autophagy^34^. Many viruses have thus evolved mechanisms to maintain mTORC1 activity during infection^35^. Recent work has shown that the LAMTOR1/2 proteins may also play important roles in xenophagy^36^. Based on these observations, SARS-CoV-2 ORF3B could play an important role in regulating mTORC1 activity and/or in the disruption of antiviral immune function.

In addition to ER/Golgi proteins, SARS-CoV-2 NSP3 interacts with the cytoplasmic RNA binding proteins FXR1 and FXR2^37^. The FXR proteins were identified as host cell components of (+)RNA Equine Encephalitis Virus (EEV) RNA replication complexes (RC)^38-40^. FXRs are recruited to the viral RC by EEV nsP3 proteins, and the FXRs are required for RC assembly. It will be interesting to determine whether the FXRs play similar roles in SARS-CoV-2 RNA replication.

## Discussion

We and others have previously reported that orthogonal PPI discovery approaches such as proximity-dependent biotinylation (BioID) can provide information that is highly complementary to IP-MS datasets^16, 41-44^. To this end, we applied BioID to identify proximity partners for SARS-CoV-2 proteins in the proteomics “workhorse” 293 cell system. This mapping project significantly expands upon the SARS-CoV-2 virus-host interactome, providing a rich resource that can be mined by the scientific community for better understanding SARS-CoV-2 pathobiology, and identifying virus-host membrane protein interactions that could be targeted in COVID-19 therapeutics.

The design and implementation of sensitive mass spectrometric approaches for the analysis of complex biological samples will be important for clinical and basic research proteomics focused on COVID-19. To this end, we also undertook an analysis of SARS-CoV-2 virions and infected Vero cell lsyates using data-dependent acquisition tandem mass spectrometry, and identified 189 unique tryptic peptides, assigned to 17 different virus proteins. This work provides a significantly expanded SARS-CoV-2 tryptic peptide compendium for use in targeted proteomics approaches such as parallel or selected reaction monitoring (PRM/SRM), or for use in spectral library building.

## Acknowledgements

FPR is supported by a Canadian Institutes of Health Research (CIHR) Foundation Grant and the Canada Excellence Research Chairs Program. Work in the A.-C.G lab is supported by a CIHR Foundation Grant, and a grant from the Natural Sciences and Engineering Research Council of Canada. A.-C.G. holds the Canada Research Chair (Tier 1) in Functional Proteomics. Work in the BR lab is supported by the Princess Margaret Cancer Foundation, and this project was generously funded by a Fast Grant from the Thistledown Foundation (Canada).

## Methods

### Cloning, Cell culture, Transfection

Individual SARS-CoV-2 ORF sequences^15^ were cloned into a Gateway compatible BioID vector, and transfected into human Flp-In T-REx 293 cells (Invitrogen, Carlsbad, CA). Following hygromycin selection and cell expansion, protein expression was induced by the addition of 1ug/ml tetracycline and 50uM biotin to culture medium (DMEM Gibson, Waltham, MA, 10% fetal calf serum, 10mM HEPES (pH 8.0) and 1% penicillin-streptomycin), as in^16^.

### SARS-CoV-2 Production

Vero E6 (African green monkey, renal epithelial) cells were maintained at 37°C, 5% CO2 in DMEM supplemented with 10% fetal bovine serum (FBS) and 1x penicillin-streptomycin. Infection with SARS-CoV-2 (TCID50=3.5×10^6^; SB3-TYAGNC isolate^6^) was performed in T175 flasks with an MOI of 0.1 in 5 mL of serum-free DMEM for 1 hour, after which infection media was replaced with 30 mL of DMEM containing 2% FBS. Culture media was collected after 48 hours.

### Sample Preparation for LC-MS Analysis of SARS-CoV-2 Proteome

Virus-containing media (TCID_50_ =1.4⨯10^7^) from Vero E6 infection was centrifuged (2 hours, 20,000 x g, 4°C) to pellet viral particles. Supernatant was removed and pellets were rinsed with cold PBS, followed by an additional centrifugation (2 hours, 20,000 x g, 4°C). PBS was removed and the pellet was lysed with 5% SDS/1% Triton X-100 in 50 mM TEAB with endonuclease (TurboNuclease, BioVision, Milpitas, CA). For proteome analysis of SARS-CoV-2 in cells, Vero E6 were infected at a MOI of 0.1, harvested 48 hours post-infection and rinsed with PBS prior to lysis. Samples were sonicated and cleared by centrifugation (10 minutes, 8,000 x g, 4°C). Cleared lysates were reduced (5 mM DTT, 60° C, 20 minutes), alkylated (10 mM iodoacetamide, RT, protected from light, 30 minutes) and acidified with H3PO4 (1.2% final concentration). Samples were diluted 1:6 with 90% methanol in 100 mM TEAB and passed through S-Trap(tm) columns (S-Trap(tm) micro kit, Protifi, Farmingdale, NY). Columns were washed with 90% methanol/100 mM TEAB 10 times prior to on-column digestion with 1 ug Trypsin/Lys-C in 50 mM TEAB (47° C, 2 hours; Cat. #V5073, Promega, Madison, WI). The resulting peptides were eluted with sequential washes of 50 mM TEAB, 0.2% HCOOH and 50% ACN/0.2% HCOOH. Eluates were lyophilized by vacuum centrifuge, reconstituted in 0.1% HCOOH, desalted with C18 columns and lyophilized prior to final reconstitution in 0.1% HCOOH for LC-MS analysis.

BioID, mass spectrometry and data analysis were conducted exactly as in^16^. For SAINT^45^ analysis, 18 MS runs from cells expressing the BirA* protein alone were used as controls. Data were searched against human RefSeq version 45 (36113 entries) supplemented with the SARS-CoV-2 proteome sequence (2019-nCoC HKU-SZ-005b^1^). For each virus bait protein, two biological replicates were each subjected to two MS runs (two technical replicates), for a total of 56 runs.

### GO enrichment

Gene ontology enrichment calculated using PANTHER (http://www.pantherdb.org/) *Homo sapiens* GO Slim dataset.

### Membrane protein prediction

Putative SARS-CoV-2 membrane spanning regions were assigned according to the TMHMM 2.0 algorithm (cbs.dtu.dk/services/TMHMM).

## Supplementary Tables

**Supp Table 1**. Virus peptide identification dataset, complete raw data. Data for viral preparation and infected cells is presented in individual tabs.

**Supp Table 2**. Complete BioID interactome data for 14 SARS-CoV-2 proteins stably expressed in 293 Flp-In T-REx cells. All high confidence PPIs are presented in Tab 1; individual bait interactomes presented in individual tabs, as indicated.

**Supp Table 3**. Complete GO enrichment data. GO enrichment for individual baits in individual tabs, as indicated.

## References

1. Chan, J.F., Kok, K.H., Zhu, Z., Chu, H., To, K.K., Yuan, S. & Yuen, K.Y. Genomic characterization of the 2019 novel human-pathogenic coronavirus isolated from a patient with atypical pneumonia after visiting Wuhan. Emerg Microbes Infect 9, 221–236 (2020).

2. Wu, F., Zhao, S., Yu, B., Chen, Y.M., Wang, W., Song, Z.G., Hu, Y., Tao, Z.W., Tian, J.H., Pei, Y.Y., Yuan, M.L., Zhang, Y.L., Dai, F.H., Liu, Y., Wang, Q.M., Zheng, J.J., Xu, L., Holmes, E.C. & Zhang, Y.Z. A new coronavirus associated with human respiratory disease in China. Nature 579, 265–269 (2020).

3. Gordon, D.E., Jang, G.M., Bouhaddou, M., Xu, J., Obernier, K., White, K.M., O’Meara, M.J., Rezelj, V.V., Guo, J.Z., Swaney, D.L., Tummino, T.A., Huttenhain, R., Kaake, R.M., Richards, A.L., Tutuncuoglu, B., Foussard, H., Batra, J., Haas, K., Modak, M., Kim, M., Haas, P., Polacco, B.J., Braberg, H., Fabius, J.M., Eckhardt, M., Soucheray, M., Bennett, M.J., Cakir, M., McGregor, M.J., Li, Q., Meyer, B., Roesch, F., Vallet, T., Mac Kain, A., Miorin, L., Moreno, E., Naing, Z.Z.C., Zhou, Y., Peng, S., Shi, Y., Zhang, Z., Shen, W., Kirby, I.T., Melnyk, J.E., Chorba, J.S., Lou, K., Dai, S.A., Barrio-Hernandez, I., Memon, D., Hernandez-Armenta, C., Lyu, J., Mathy, C.J.P., Perica, T., Pilla, K.B., Ganesan, S.J., Saltzberg, D.J., Rakesh, R., Liu, X., Rosenthal, S.B., Calviello, L., Venkataramanan, S., Liboy-Lugo, J., Lin, Y., Huang, X.P., Liu, Y., Wankowicz, S.A., Bohn, M., Safari, M., Ugur, F.S., Koh, C., Savar, N.S., Tran, Q.D., Shengjuler, D., Fletcher, S.J., O’Neal, M.C., Cai, Y., Chang, J.C.J., Broadhurst, D.J., Klippsten, S., Sharp, P.P., Wenzell, N.A., Kuzuoglu-Ozturk, D., Wang, H.Y., Trenker, R., Young, J.M., Cavero, D.A., Hiatt, J., Roth, T.L., Rathore, U., Subramanian, A., Noack, J., Hubert, M., Stroud, R.M., Frankel, A.D., Rosenberg, O.S., Verba, K.A., Agard, D.A., Ott, M., Emerman, M., Jura, N., von Zastrow, M., Verdin, E., Ashworth, A., Schwartz, O., d’Enfert, C., Mukherjee, S., Jacobson, M., Malik, H.S., Fujimori, D.G., Ideker, T., Craik, C.S., Floor, S.N., Fraser, J.S., Gross, J.D., Sali, A., Roth, B.L., Ruggero, D., Taunton, J., Kortemme, T., Beltrao, P., Vignuzzi, M., Garcia-Sastre, A., Shokat, K.M., Shoichet, B.K. & Krogan, N.J. A SARS-CoV-2 protein interaction map reveals targets for drug repurposing. Nature 583, 459–468 (2020).

4. Stukalov, A., Girault, V., Grass, V., Bergant, V., Karayel, O., Urban, C., Haas, D.A., Huang, Y., Oubraham, L., Wang, A., Hamad, S.M., Piras, A., Tanzer, M., Hansen, F.M., Enghleitner, T., Reinecke, M., Lavacca, T.M., Ehmann, R., Wölfel, R., Jores, J., Kuster, B., Protzer, U., Rad, R., Ziebuhr, J., Thiel, V., Scaturro, P., Mann, M. & Pichlmair, A. Multi-level proteomics reveals host-perturbation strategies of SARS-CoV-2 and SARS-CoV. bioRxiv (2020).

5. Roux, K.J., Kim, D.I., Raida, M. & Burke, B. A promiscuous biotin ligase fusion protein identifies proximal and interacting proteins in mammalian cells. J Cell Biol 196, 801–810 (2012).

6. Banerjee, A., Nasir, J.A., Budylowski, P., Yip, L., Aftanas, P., Christie, N., Ghalami, A., Baid, K., Raphenya, A.R., Hirota, J.A., Miller, M.S., McGeer, A.J., Ostrowski, M., Kozak, R.A., McArthur, A.G., Mossman, K. & Mubareka, S. Isolation, Sequence, Infectivity, and Replication Kinetics of Severe Acute Respiratory Syndrome Coronavirus 2. Emerg Infect Dis 26, 2054–2063 (2020).

7. Zecha, J., Lee, C.Y., Bayer, F.P., Meng, C., Grass, V., Zerweck, J., Schnatbaum, K., Michler, T., Pichlmair, A., Ludwig, C. & Kuster, B. Data, reagents, assays and merits of proteomics for SARS-CoV-2 research and testing. Mol Cell Proteomics (2020).

8. Gouveia, D., Grenga, L., Gaillard, J.C., Gallais, F., Bellanger, L., Pible, O. & Armengaud, J. Shortlisting SARS-CoV-2 Peptides for Targeted Studies from Experimental Data-Dependent Acquisition Tandem Mass Spectrometry Data. Proteomics 20, e2000107 (2020).

9. Grenga, L., Gallais, F., Pible, O., Gaillard, J.C., Gouveia, D., Batina, H., Bazaline, N., Ruat, S., Culotta, K., Miotello, G., Debroas, S., Roncato, M.A., Steinmetz, G., Foissard, C., Desplan, A., Alpha-Bazin, B., Almunia, C., Gas, F., Bellanger, L. & Armengaud, J. Shotgun proteomics analysis of SARS-CoV-2-infected cells and how it can optimize whole viral particle antigen production for vaccines. Emerg Microbes Infect 9, 1712–1721 (2020).

10. Liang, J.R., Lingeman, E., Luong, T., Ahmed, S., Muhar, M., Nguyen, T., Olzmann, J.A. & Corn, J.E. A Genome-wide ER-phagy Screen Highlights Key Roles of Mitochondrial Metabolism and ER-Resident UFMylation. Cell 180, 1160–1177 e1120 (2020).

11. Bojkova, D., Klann, K., Koch, B., Widera, M., Krause, D., Ciesek, S., Cinatl, J. & Munch, C. Proteomics of SARS-CoV-2-infected host cells reveals therapy targets. Nature 583, 469–472 (2020).

12. Ihling, C., Tanzler, D., Hagemann, S., Kehlen, A., Huttelmaier, S., Arlt, C. & Sinz, A. Mass Spectrometric Identification of SARS-CoV-2 Proteins from Gargle Solution Samples of COVID-19 Patients. J Proteome Res (2020).

13. Nikolaev, E.N., Indeykina, M.I., Brzhozovskiy, A.G., Bugrova, A.E., Kononikhin, A.S., Starodubtseva, N.L., Petrotchenko, E.V., Kovalev, G.I., Borchers, C.H. & Sukhikh, G.T. Mass-Spectrometric Detection of SARS-CoV-2 Virus in Scrapings of the Epithelium of the Nasopharynx of Infected Patients via Nucleocapsid N Protein. J Proteome Res (2020).

14. Gouveia, D., Miotello, G., Gallais, F., Gaillard, J.C., Debroas, S., Bellanger, L., Lavigne, J.P., Sotto, A., Grenga, L., Pible, O. & Armengaud, J. Proteotyping SARS-CoV-2 Virus from Nasopharyngeal Swabs: A Proof-of-Concept Focused on a 3 Min Mass Spectrometry Window. J Proteome Res (2020).

15. Kim, D.K., Knapp, J.J., Kuang, D., Chawla, A., Cassonnet, P., Lee, H., Sheykhkarimli, D., Samavarchi-Tehrani, P., Abdouni, H., Rayhan, A., Li, R., Pogoutse, O., Coyaud, E., van der Werf, S., Demeret, C., Gingras, A.C., Taipale, M., Raught, B., Jacob, Y. & Roth, F.P. A Comprehensive, Flexible Collection of SARS-CoV-2 Coding Regions. G3 (Bethesda) (2020).

16. Coyaud, E., Ranadheera, C., Cheng, D.T., Goncalves, J., Dyakov, B., Laurent, E., St-Germain, J.R., Pelletier, L., Gingras, A.C., Brumell, J.H., Kim, P.K., Safronetz, D. & Raught, B. Global interactomics uncovers extensive organellar targeting by Zika virus. Mol Cell Proteomics (2018).

17. Knight, J.D.R., Choi, H., Gupta, G.D., Pelletier, L., Raught, B., Nesvizhskii, A.I. & Gingras, A.C. ProHits-viz: a suite of web tools for visualizing interaction proteomics data. Nat Methods 14, 645–646 (2017).

18. Chitwood, P.J. & Hegde, R.S. An intramembrane chaperone complex facilitates membrane protein biogenesis. Nature 584, 630–634 (2020).

19. Wu, H., Carvalho, P. & Voeltz, G.K. Here, there, and everywhere: The importance of ER membrane contact sites. Science 361 (2018).

20. Belov, G.A. & van Kuppeveld, F.J. (+)RNA viruses rewire cellular pathways to build replication organelles. Curr Opin Virol 2, 740–747 (2012).

21. Roulin, P.S., Lotzerich, M., Torta, F., Tanner, L.B., van Kuppeveld, F.J., Wenk, M.R. & Greber, U.F. Rhinovirus uses a phosphatidylinositol 4-phosphate/cholesterol counter-current for the formation of replication compartments at the ER-Golgi interface. Cell Host Microbe 16, 677–690 (2014).

22. van der Schaar, H.M., Dorobantu, C.M., Albulescu, L., Strating, J. & van Kuppeveld, F.J.M. Fat(al) attraction: Picornaviruses Usurp Lipid Transfer at Membrane Contact Sites to Create Replication Organelles. Trends Microbiol 24, 535–546 (2016).

23. Zhang, Z., He, G., Filipowicz, N.A., Randall, G., Belov, G.A., Kopek, B.G. & Wang, X. Host Lipids in Positive-Strand RNA Virus Genome Replication. Front Microbiol 10, 286 (2019).

24. Antonny, B., Bigay, J. & Mesmin, B. The Oxysterol-Binding Protein Cycle: Burning Off PI(4)P to Transport Cholesterol. Annu Rev Biochem 87, 809–837 (2018).

25. Stoeck, I.K., Lee, J.Y., Tabata, K., Romero-Brey, I., Paul, D., Schult, P., Lohmann, V., Kaderali, L. & Bartenschlager, R. Hepatitis C Virus Replication Depends on Endosomal Cholesterol Homeostasis. J Virol 92 (2018).

26. Carmona-Gutierrez, D., Bauer, M.A., Zimmermann, A., Kainz, K., Hofer, S.J., Kroemer, G. & Madeo, F. Digesting the crisis: autophagy and coronaviruses. Microb Cell 7, 119–128 (2020).

27. English, L., Chemali, M., Duron, J., Rondeau, C., Laplante, A., Gingras, D., Alexander, D., Leib, D., Norbury, C., Lippe, R. & Desjardins, M. Autophagy enhances the presentation of endogenous viral antigens on MHC class I molecules during HSV-1 infection. Nat Immunol 10, 480–487 (2009).

28. Wong, H.H. & Sanyal, S. Manipulation of autophagy by (+) RNA viruses. Semin Cell Dev Biol 101, 3–11 (2020).

29. Ren, Z., Ding, T., Zuo, Z., Xu, Z., Deng, J. & Wei, Z. Regulation of MAVS Expression and Signaling Function in the Antiviral Innate Immune Response. Front Immunol 11, 1030 (2020).

30. Shi, C.S., Qi, H.Y., Boularan, C., Huang, N.N., Abu-Asab, M., Shelhamer, J.H. & Kehrl, J.H. SARS-coronavirus open reading frame-9b suppresses innate immunity by targeting mitochondria and the MAVS/TRAF3/TRAF6 signalosome. J Immunol 193, 3080–3089 (2014).

31. Jiang, H.W., Zhang, H.N., Meng, Q.F., Xie, J., Li, Y., Chen, H., Zheng, Y.X., Wang, X.N., Qi, H., Zhang, J., Wang, P.H., Han, Z.G. & Tao, S.C. SARS-CoV-2 Orf9b suppresses type I interferon responses by targeting TOM70. Cell Mol Immunol (2020).

32. Yarbrough, M.L., Mata, M.A., Sakthivel, R. & Fontoura, B.M. Viral subversion of nucleocytoplasmic trafficking. Traffic 15, 127–140 (2014).

33. Yuen, C.K., Lam, J.Y., Wong, W.M., Mak, L.F., Wang, X., Chu, H., Cai, J.P., Jin, D.Y., To, K.K., Chan, J.F., Yuen, K.Y. & Kok, K.H. SARS-CoV-2 nsp13, nsp14, nsp15 and orf6 function as potent interferon antagonists. Emerg Microbes Infect 9, 1418–1428 (2020).

34. Condon, K.J. & Sabatini, D.M. Nutrient regulation of mTORC1 at a glance. J Cell Sci 132 (2019).

35. Le Sage, V., Cinti, A., Amorim, R. & Mouland, A.J. Adapting the Stress Response: Viral Subversion of the mTOR Signaling Pathway. Viruses 8 (2016).

36. Lin, C.Y., Nozawa, T., Minowa-Nozawa, A., Toh, H., Aikawa, C. & Nakagawa, I. LAMTOR2/LAMTOR1 complex is required for TAX1BP1-mediated xenophagy. Cell Microbiol 21, e12981 (2019).

37. Li, Y. & Zhao, X. Concise review: Fragile X proteins in stem cell maintenance and differentiation. Stem Cells 32, 1724–1733 (2014).

38. Meshram, C.D., Phillips, A.T., Lukash, T., Shiliaev, N., Frolova, E.I. & Frolov, I. Mutations in Hypervariable Domain of Venezuelan Equine Encephalitis Virus nsP3 Protein Differentially Affect Viral Replication. J Virol 94 (2020).

39. Frolov, I., Kim, D.Y., Akhrymuk, M., Mobley, J.A. & Frolova, E.I. Hypervariable Domain of Eastern Equine Encephalitis Virus nsP3 Redundantly Utilizes Multiple Cellular Proteins for Replication Complex Assembly. J Virol 91 (2017).

40. Kim, D.Y., Reynaud, J.M., Rasalouskaya, A., Akhrymuk, I., Mobley, J.A., Frolov, I. & Frolova, E.I. New World and Old World Alphaviruses Have Evolved to Exploit Different Components of Stress Granules, FXR and G3BP Proteins, for Assembly of Viral Replication Complexes. PLoS Pathog 12, e1005810 (2016).

41. Coyaud, E., Mis, M., Laurent, E.M., Dunham, W.H., Couzens, A.L., Robitaille, M., Gingras, A.C., Angers, S. & Raught, B. BioID-based Identification of Skp Cullin F-box (SCF)beta-TrCP1/2 E3 Ligase Substrates. Mol Cell Proteomics 14, 1781–1795 (2015).

42. Gingras, A.C., Abe, K.T. & Raught, B. Getting to know the neighborhood: using proximity-dependent biotinylation to characterize protein complexes and map organelles. Curr Opin Chem Biol 48, 44–54 (2019).

43. Hesketh, G.G., Youn, J.Y., Samavarchi-Tehrani, P., Raught, B. & Gingras, A.C. Parallel Exploration of Interaction Space by BioID and Affinity Purification Coupled to Mass Spectrometry. Methods Mol Biol 1550, 115–136 (2017).

44. Lambert, J.P., Tucholska, M., Go, C., Knight, J.D. & Gingras, A.C. Proximity biotinylation and affinity purification are complementary approaches for the interactome mapping of chromatin-associated protein complexes. J Proteomics 118, 81–94 (2015).

45. Choi, H., Larsen, B., Lin, Z.Y., Breitkreutz, A., Mellacheruvu, D., Fermin, D., Qin, Z.S., Tyers, M., Gingras, A.C. & Nesvizhskii, A.I. SAINT: probabilistic scoring of affinity purification-mass spectrometry data. Nat Methods 8, 70–73 (2011).

